# Combined Checkpoint Inhibition Amplifies Post-Infarction Injury via T Cell-Mediated Macrophage Activation

**DOI:** 10.64898/2026.05.18.726115

**Authors:** Xingzhi Wang, Miaomiao Cai, Yong Zhou, Mengyuan Feng, Peibin Chen, Junneng Zhang, Siqi Liu, Yajie Song, Cuige Zhu, Ailan Chen, Guoshuai Feng

## Abstract

**Background:** This study aimed to investigate whether combined PD-1/CTLA-4 immune checkpoint inhibition predisposes the heart to a hyperinflammatory state, thereby exacerbating cardiac injury following acute myocardial infarction (MI), a critical unresolved question in cardio-oncology.

**Methods:** Myocardial infarction was induced in *Pd1*^−/−^*Ctla4*^+/−^ mice, a genetic model mimicking combined checkpoint inhibition. Key mechanistic insights were gained through *in vivo*depletion of CD8^+^ T cells (using anti-CD8a antibody) and pharmacological inhibition of the JAK-STAT1 pathway (using Tofacitinib). Cardiac function, structural injury, and immune responses were comprehensively assessed via echocardiography, flow cytometry, immunofluorescence, and molecular analyses.

**Results:** Compared to wild-type controls, *Pd1*^−/−^*Ctla4*^+/−^ mice exhibited significantly increased post-MI mortality, worse cardiac function, and larger infarct size. Mechanistically, the aggravated injury was driven by an amplified infiltration of activated, IFN-γ-producing CD8^+^ T cells, which activated the JAK-STAT1 pathway in macrophages, polarizing them towards a pro-inflammatory state. Depleting CD8^+^ T cells or inhibiting the JAK-STAT1 pathway effectively attenuated macrophage-driven inflammation and improved all aspects of post-MI injury.

**Conclusions:** Combined PD-1/CTLA-4 blockade exacerbates post-infarction cardiac injury by promoting CD8^+^ T cell-mediated activation of macrophages via the JAK-STAT1 axis. This work elucidates MI as a context-dependent immune-related adverse event in ICI therapy and identifies CD8^+^ T cells and the JAK-STAT1 pathway as promising therapeutic targets for cardioprotection in these patients.

**RESEARCH PERSPECTIVE:** *What Is New?:*

- This study identifies acute myocardial infarction (MI) as a potential, context-dependent immune-related adverse event in the setting of combined PD-1/CTLA-4 checkpoint inhibition, shifting the paradigm beyond the classic focus on myocarditis.
- It elucidates a novel pathogenic axis where combined checkpoint deficiency exacerbates post-MI injury specifically through CD8^+^ T cell-derived IFN-γ, which activates macrophages via the JAK-STAT1 pathway.

*What Question Should Be Addressed Next?:*

- Future studies should employ anti-PD-1/CTLA-4 monoclonal antibodies in wild-type or humanized mouse models to validate findings and better recapitulate the pharmacokinetics of clinical ICI therapy, strengthening translational relevance.
- The long-term consequences of this primed inflammatory state on chronic cardiac remodeling, heart failure development, and the potential interplay with atherosclerosis warrant further investigation.

## Introduction

Immune checkpoint inhibitors (ICIs) have markedly improved outcomes in patients with advanced cancers^[1]^. However, their use is associated with immune-related adverse events (IRAEs) due to systemic immune overactivation^[2, 3]^. Among these, ICI-induced myocarditis represents one of the most lethal IRAEs, albeit with a relatively low frequency. Even in patients without clinically apparent myocarditis, a key unanswered question is whether ICIs alter cardiac immunomodulation^[4–6]^. Specifically, whether ICIs elevate the baseline inflammatory tone or prime the heart toward a subcritical, hyper-inflammatory state^[7]^. If so, such priming could predispose the heart to an exaggerated inflammatory response and more severe injury following an acute cardiac insult.

Acute myocardial infarction (MI) is a common and catastrophic cardiac emergency. The cardiovascular risk of administering ICIs to patients with pre-existing heart disease remains poorly understood^[8–10]^. In particular, it is unclear whether an acute MI during ICI treatment triggers a more intense inflammatory reaction and consequently leads to greater cardiac damage. Thus, investigating whether ICIs exacerbate post-MI injury is of significant clinical importance^[11–14]^.

Currently approved ICIs primarily target *Pd1*, PDL1, *Ctla4*, and LAG3^[15–22]^. Among these, combined *Pd1* and *Ctla4* inhibition has emerged as a pivotal therapeutic strategy to enhance antitumor immunity^[23–25]^. However, this combination also increases the risk of ICI-associated myocarditis^[26–28]^. Whether it influences subclinical myocardial inflammation and aggravates cardiac injury after acute MI requires further investigation^[29–31]^.

This study aimed to determine whether combined blockade of *Pd1* and *Ctla4* promotes a mild inflammatory state in the heart, thereby exacerbating myocardial injury after infarction. We employed a genetic model (*Pd1*^−/−^*Ctla4*^+/−^ mice), established in our prior work, to mimic the systemic immunologic effects of dual checkpoint inhibition^[32]^. To specifically assess the impact of this preconditioned inflammatory state on post-infarction remodeling, we screened *Pd1*^−/−^*Ctla4*^+/−^ mice by echocardiography prior to surgery and included only those without baseline signs of myocarditis in the myocardial infarction study. Cardiac remodeling was then comprehensively assessed through survival analysis, functional measurements, and histopathological examination. Compared with *Pd1*^*+/+*^*Ctla4*^*+/+*^ controls, *Pd1*^−/−^*Ctla4*^+/−^ mice exhibited significantly exacerbated post-MI injury, including reduced survival, worse cardiac function, larger infarct size, and aggravated myocardial hypertrophy, fibrosis, inflammation, and apoptosis. Increased infiltration of activated CD8^+^ T cells and CD64^+^ macrophages was observed in the hearts of deficient mice. RNA sequencing revealed marked activation of the JAK-STAT1 pathway and upregulation of inflammatory cytokines such as IFN-γ. In vivo depletion of CD8^+^ T cells or pharmacological inhibition of the JAK-STAT1 pathway attenuated macrophage-driven inflammation, improved cardiac function, and ameliorated hypertrophy, fibrosis, and apoptosis after MI.

In summary, this study demonstrates that combined *Pd1* and *Ctla4* inhibition exacerbates post-MI cardiac injury by promoting CD8^+^ T cell-mediated aggravation of macrophage inflammation via the JAK-STAT1 axis. These findings confirm that dual checkpoint inhibition primes the heart toward a hyper-inflammatory state, leading to an exaggerated response and more severe injury after acute MI. CD8^+^ T cells and the JAK-STAT1 pathway represent two promising therapeutic targets for mitigating such cardiac adverse events in patients receiving combination ICI therapy.

## METHODS

### Data Availability Statement

Data are available in the article and supplementary material. Raw microscopy images will be shared upon reasonable request to the corresponding author.

### Mice

All experiments were conducted using male or female mice aged 12 weeks. All mice strains were purchased from the generated by Shanghai Model Organisms Center, Inc. All mice were maintained at the Laboratory Animal Center of Guangzhou Medical University (Guangzhou, China) under standard laboratory conditions with food and water provided ad libitum. All animal care and handling procedures were conducted in accordance with protocols approved by the Ethics Committee of Guangzhou Medical University (protocol no. GY2024-365).

### Animal Studies

Animal procedures were approved by the Institutional Animal Care and Use Committee at Guangzhou Medical University following the recommendations in the Guide for the Care and Use of Laboratory Animals of the NIH guidelines. *Pd1* global knockout mice and *Ctla4* heterozygous knockout mouse were purchased from Shanghai Model Organisms Center, Inc. (Shanghai, China). All mouse genotypes were confirmed according to established protocols.

### myocardial infarction (MI) injury and antibody injection

For myocardial infarction injury, mice were anesthetized with sodium pentobarbital, intubated, and mechanically ventilated^[33]^. A left thoracotomy was performed through the third to fourth intercostal space, followed by blunt dissection of the pectoralis major and minor muscles to expose the heart. The left anterior descending coronary artery was ligated using 8-0 prolene sutures with needles^[34]^. The thoracic cavity was closed sequentially with 5-0 sutures for ribs, muscles, and skin. Eye ointment was applied to prevent corneal dehydration during surgery. Postoperatively, mice were recovered on a heated pad, resuscitated with normal saline, and received gentamicin for infection prophylaxis. Control animals underwent identical procedures except the coronary ligature was tied as a loose knot (sham ligation)^[35]^. Health *Pd1*^*−/−*^*Ctla4*^*−/−*^ mice were intraperitoneally administered 200 µL of anti-CD8 antibody (BioXCell, Cat. BE0061) or isotype control every other day^[36–38]^. The treatment continued until 27 days after surgery.

### Echocardiography

Mouse echocardiography was performed in Guangzhou Medical University Scientific Research Center using the Visual Sonics Vevo 2100 Echocardiography System. 2D and M-mode images were obtained in the long and short axis views. Ejection fraction (EF) and LV dimensions were calculated using Vevo LAB software and standard techniques. Measurements were performed on 3 independently acquired images per animal, by investigators who were blinded to experimental group. Each experimental group included at least 6 animals.

### Immunofluorescence staining

For murine fresh tissues, fixation was performed with 4% paraformaldehyde, followed by dehydration in 30% sucrose solution and embedding in OCT compound. The sections were blocked with 5% BSA blocking buffer, and incubated with primary antibodies overnight at 4°C, including: CD8a Monoclonal Antibody (Cell Signaling, cat# 98941), Granzyme B Monoclonal Antibody (Cell Signaling, cat# 17215), CD3 Monoclonal Antibody (Cell Signaling, cat# 73484), Anti-Interferon gamma antibody (Abcam, cat# EPR28352-7)^[39]^. After washing with TBST, sections were incubated with corresponding secondary antibodies at room temperature for 2 h. Fluorescence signals were imaged using a Zeiss confocal microscope.

### Terminal deoxynucleotidyl transferase dUTP Nick-End Labeling (TUNEL) staining

Paraffin-embedded sections were deparaffinized, rehydrated, and digested with proteinase K. The TUNEL reaction mixture, containing terminal deoxynucleotidyl transferase (TdT) and fluorescein-labeled dUTP, was applied to cover the tissue sections, followed by incubation at 37 °C in the dark for 60 min. After thorough washing, fluorescent signals were visualized using a Zeiss confocal microscope.

### Trichrome staining and wheat germ agglutinin (WGA) staining

Mouse hearts were fixed overnight in 4% paraformaldehyde at 4°C, dehydrated through an ethanol series, cleared with xylene, embedded in paraffin, and sectioned at 6 μm thickness. Sections were deparaffinized, rehydrated, and subjected to Masson’s trichrome staining for cardiac fibrosis assessment, with quantitative analysis performed using Image J software^[34, 40]^. For cardiomyocyte cross-sectional area measurement, paraffin sections were stained with rhodamine-conjugated wheat germ agglutinin (WGA), visualized using a Zeiss confocal microscope, and analyzed with Zeiss Axiovision software.

### Bone-marrow derived macrophages (BMDMs) generation and stimulation

To generate bone marrow-derived macrophages (BMDM), bone marrow cells isolated from the femurs and tibias of mice were cultured in DMEM supplemented with 10% fetal bovine serum (FBS), 20 ng/mL macrophage colony-stimulating factor (M-CSF), 1% penicillin, and 1% streptomycin for 7 days^[41, 42]^. On day 7, the medium was replaced with M-CSF-free medium for an additional 2 days^[43]^. Subsequently, 1×10^6^ BMDMs were seeded into 6-well plates and cultured overnight to allow for adherence, followed by stimulation for real-time quantitative PCR and Western blot analysis. On day 8, the medium was replaced with M-CSF-free medium. For qPCR and Western blot assays following treatment with JAK-STAT1 inhibitors, BMDMs were seeded in 6-well plates and cultured overnight to allow adhesion.

### RT-PCR

Total RNA was extracted from cardiac tissues or cells using a commercial RNA extraction kit. The concentration and purity of RNA were measured using a NanoDrop spectrophotometer. RNA was then reverse-transcribed into complementary DNA (cDNA). Quantitative real-time PCR (qPCR) was performed with sequence-specific primers using SYBR Green master mix in a 20 μl reaction system. mRNA expression levels were normalized to the housekeeping gene Gapdh. The sequences of primers used are listed in Supplemental Table 4.

### Western blot

Mouse cardiac tissues and cells were collected and homogenized in RIPA lysis buffer containing protease inhibitors. Total protein concentration was measured using the BCA Protein Assay Kit. Primary antibodies included: Anti-PD1 antibody (Abcam, cat# ab214421), Ctla4 Rabbit Monoclonal Antibody (Cell Signaling, cat# 53560), Anti-Interferon gamma antibody (Abcam, cat# EPR28352-7), Phospho-Stat1 (Tyr701) Rabbit Monoclonal Antibody (Cell Signaling, cat# 9167), TNF-alpha Rabbit Monoclonal Antibody (Cell Signaling, cat# 11948), IL-6 Rabbit Monoclonal Antibody (Cell Signaling, cat# 12912), IL-1β Mouse mAb (Cell Signaling, cat# 12242), α-Tubulin Antibody (Cell Signaling, cat# 2144), Caspase-3 Antibody (Cell Signaling, cat# 9662), and iNOS Rabbit Monoclonal Antibody (Cell Signaling, cat# 13120). The membranes were incubated with primary antibodies overnight at 4°C, followed by detection with horseradish peroxidase-conjugated secondary antibodies. Immunoreactive bands were visualized using a chemiluminescent substrate. Band intensity was quantified with ImageJ software and normalized to α-Tubulin.

### Flow cytometry

Mouse hearts were harvested after saline perfusion, minced, and digested with collagenase I, hyaluronidase, and DNase I at 37°C for 30 minutes. Digestion was terminated by adding PBS supplemented with 2% fetal bovine serum and 0.2% bovine serum albumin. The cell suspension was filtered through a 40 μm cell strainer. Red blood cells were lysed with ACK lysis buffer, and the remaining cells were resuspended in PBS containing 2% FBS and 2 mM EDTA for antibody staining at 4°C for 30 minutes.

Single-cell suspensions were prepared according to standard flow cytometry protocols, followed by sequential antibody staining. Cell sorting was performed using the BD FACS Aria III flow cytometer, and the following gating strategy was applied: debris was excluded via an FSC-A/SSC-A gate, doublets were excluded through an FSC-H/FSC-W gate, and target cell populations were finally identified by fluorescence gating. Throughout the sorting procedure, cells were maintained on ice or at 4°C. Sorted cells were collected in tubes containing high-protein medium to preserve cell viability.

### Statistical analysis

All experimental data are presented as mean ± standard deviation (SD). Statistical analyses were performed using GraphPad Prism software (version 10). For comparisons between two groups, the unpaired two-tailed Student’s t-test was used for data conforming to a normal distribution; Welch’s correction was applied when variances were unequal. For data that did not meet the assumption of normality, the nonparametric Mann-Whitney U test was employed. For comparisons among multiple groups, one-way or two-way analysis of variance (ANOVA) was used as appropriate. A P value of less than 0.05 was considered statistically significant, denoted as \**P* < 0.05, \*\**P* < 0.01, and \*\*\**P* < 0.001; “ns” indicates no statistical significance.

## RESULTS

### Combined deficiency of *Pd1* and *Ctla4* exacerbates cardiac injury after myocardial infarction

To model combined PD1 and CTLA4 inhibition, we used *Pd1* and *Ctla4* gene knockout mice. In line with established literature, *Pd1*^*+/−*^, *Pd1*^*−/−*^, and *Ctla4*^*+/−*^ mice were viable and fertile, while *Ctla4*^*−/−*^ mice uniformly developed fatal systemic autoimmunity, succumbing within three weeks^[32]^. We therefore crossed *Pd1*^*−/−*^ mice with *Ctla4*^*+/−*^ mice to generate *Pd1*^*−/−*^*Ctla4*^*+/−*^ offspring, which served as the genetic model for dual checkpoint inhibition in this study (**Figure S1A-S1C**). Within the resulting *Pd1*^*−/−*^*Ctla4*^*+/−*^ cohort, approximately one-quarter of mice exhibited progressive weight loss and died within 3 months of birth. In contrast, the other three-quarters remained healthy with a normal lifespan (**Figure S1D)**. This early-mortality subset exhibited poor general condition, significantly lower body weight, and pronounced cardiac dysfunction, accompanied by elevated inflammatory markers and increased immune cell infiltration-all features consistent with severe myocarditis (**Figure S1E-S1H**).

For the subsequent MI study, we established two experimental cohorts. *Pd1*^*−/−*^*Ctla4*^*+/−*^ mice were prescreened by echocardiography, overall condition and body weight monitoring to exclude individuals with baseline cardiac dysfunction or signs of myocarditis. Mice meeting these criteria underwent MI surgery which were named Model mice. Age- and sex-matched wild-type (*Pd1*^*+/+*^*Ctla4*^*+/+*^) mice were included as the control group and subjected to the same surgical procedure (**Figure 1A; Figure S2A**). Compared with wild-type (*Pd1*^*−/−*^*Ctla4*^*+/−*^) controls, *Pd1*^*−/−*^*Ctla4*^*+/−*^ mice exhibited significantly higher post-MI mortality (**Figure 1B**). Assessment of cardiac injury revealed a larger infarct size (**Figure 1C**), increased cardiomyocyte death in the peri-infarct border zone (**Figure 1D**), and elevated expression of apoptotic signaling proteins in *Pd1*^*−/−*^*Ctla4*^*+/−*^ mice (**Figure S2B**). Furthermore, these mice developed worse cardiac function, evidenced by reduced ejection fraction and fractional shortening, accompanied by pathological left ventricular chamber dilation (**Figure 1E; Table S1**). Cardiac remodeling was also exacerbated, as indicated by aggravated cardiac hypertrophy (**Figure 1F and 1G; Figure S2C**) and enhanced interstitial fibrosis (**Figure S2D and S2E**). Collectively, these data demonstrate that combined deficiency of *Pd1* and *Ctla4* exacerbates cardiac injury and adverse remodeling after myocardial infarction.

**Figure 1:**
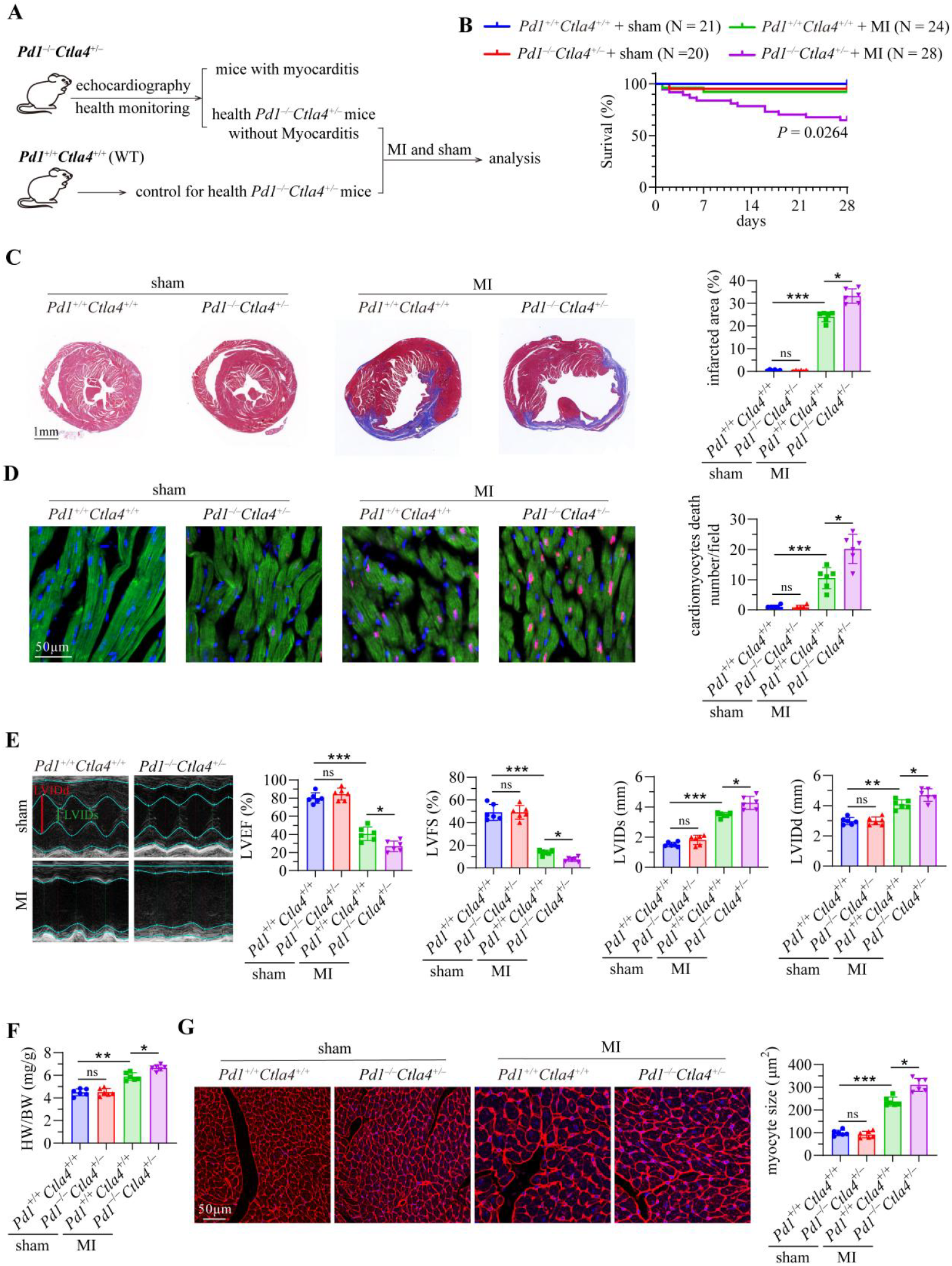
Combined deficiency of *Pd1* and *Ctla4* exacerbates cardiac injury after myocardial infarction. **A**, Schematic of the experimental strategy. Healthy *Pd1*^−/−^*Ctla4*^+/−^mice were screened via echocardiography and body weight monitoring. Mice without myocarditis were designated as the PCNM group; wild-type (*Pd1*^*+/+*^*Ctla4*^*+/+*^) mice served as the WT controls. Both groups subsequently underwent either myocardial infarction (MI) or sham surgery. **B**, Kaplan-Meier survival curves of mice for 4 weeks following MI surgery. **C**, Representative Masson’s trichrome staining images (left) and quantification of myocardial infarction area proportion (right). Collagen deposition: blue; cardiomyocytes: red; nuclei:black. **D**, Representative TUNEL staining images and quantification of apoptotic (TUNEL^+^) cardiomyocytes in the peri-infarct border zone 28 days after surgery. TUNEL^+^ nuclei are shown in red, cTnT^+^ cardiomyocytes in green, and DAPI-counterstained nuclei in blue. **E**, Representative echocardiographic images and quantification of left ventricular ejection fraction (LVEF) and fractional shortening (LVFS) 28 days post-surgery. **F**, Quantification of the heart weight to body weight (HW/BW) ratio. **G**, Representative WGA staining images (left) and quantification of cardiomyocyte size based on WGA staining (right). Data are presented as mean ± SD. ns indicates non-significance, \**P*<0.05, \*\**P*<0.01,\*\*\**P*<0.001.

### Loss of *Pd1* and *Ctla4* augments CD8^+^ T cells infiltration in the heart after MI

We examined cardiac inflammation 4 days after myocardial infarction (MI) and sham surgery in wild-type (*Pd1*^*+/+*^*Ctla4*^*+/+*^) and *Pd1*^*−/−*^*Ctla4*^*+/−*^ mice (**Figure 1A; Figure S3A; Figure S4A**). Quantitative PCR showed that MI–but not sham surgery–triggered a significant upregulation of the proinflammatory cytokine genes *Il1b,Il6, Tnf*, and*Ifng*in *Pd1*^*−/−*^*Ctla4*^*+/−*^ hearts relative to WT controls (**Figure 2B; Figure S4B**). Flow cytometry revealed a significant post-MI increase of CD8^*+*^ T cells in *Pd1*^*−/−*^*Ctla4*^*+/−*^ hearts compared to WT controls (**Figure 2C-2E**). Immunofluorescence staining demonstrated marked accumulation of CD8^+^ T cells in the hearts of knockout mice after MI, with increased expression of granzyme B (GzmB) indicating an enhanced activation state of these cells (**Figure 2F-2H**). CD64^*+*^ macrophages, Ly6G^+^ neutrophils, CD19^*+*^ B and CD4^+^ T cells showed no pronounced elevation in *Pd1*^*−/−*^*Ctla4*^*+/−*^ hearts (**Figure S3B and S3C**). In contrast, sham surgery induced no significant difference in immune cell numbers between *Pd1*^*−/−*^*Ctla4*^*+/−*^ and WT mice (**Figure S4C and S4D**). Macrophages constituted the most abundant population of infiltrating immune cells in the heart after MI. We sorted CD64^+^ macrophages and performed micropre-amplification PCR, which revealed significantly higher expression of the inflammatory cytokine genes*Il1b,Il6*, and *Tnf* in macrophages isolated from *Pd1*^*−/−*^*Ctla4*^*+/−*^ hearts compared to those from WT hearts after MI (**Figure 2I**).

**Figure 2:**
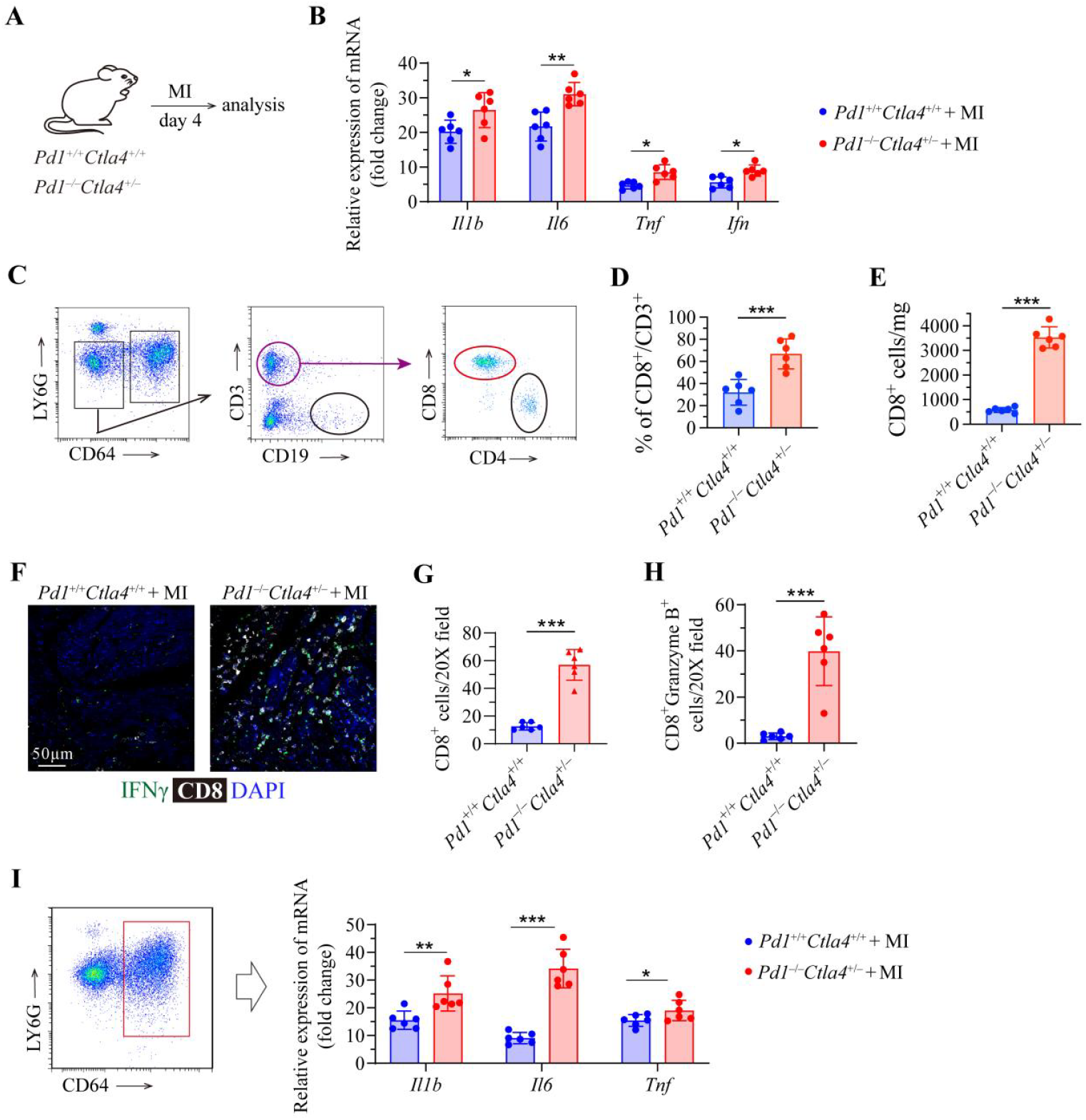
Immune cell profiling in the heart after myocardial infarction. **A**, Schematic of the experimental timeline. 12-week-old wild-type (*Pd1*^*+/+*^*Ctla4*^*+/+*^) and *Pd1*^*−/−*^*Ctla4*^*+/−*^ mice underwent permanent coronary artery ligation or sham surgery, with analysis performed 4 days later. **B**, Quantitative PCR analysis of *Il1b, Il6, Tnf*and *Ifng* mRNA levels in heart tissue. **C**, Gating strategy for the identification by flow cytometry of CD45^+^ immune cells, Ly6G^+^ neutrophils, CD19^+^ B cells, CD3^+^ T cells, and CD4^+^ or CD8^+^ T cell subsets. **D**, Quantification of CD8^+^ T cells per milligram of heart tissue by flow cytometry. **E**, Proportion of CD8^+^ T cells within the CD3^+^ T cell population. **F-H**, Representative immunofluorescence images (**F, G**) and quantitative analysis (**H**) of CD8 and granzyme B (GZMB) co-localization in heart sections. **I**, Schematic of the flow-cytometric sorting strategy for CD64^+^ macrophages (left) and quantitative PCR analysis of Il1b, Il6 and Tnf expression in sorted macrophages (right). Data are presented as mean±SD. ns, not significant; \**P* < 0.05, \*\**P*< 0.01, \*\*\**P*< 0.001.

Collectively, these findings indicate that combined deficiency of *Pd1* and *Ctla4* promotes the recruitment of activated, pro-inflammatory CD8^+^ T cells into the heart after myocardial infarction, which in turn may exacerbate cardiac inflammation by enhancing the inflammatory activity of macrophages.

### CD8^+^ T Cell Depletion ameliorates cardiac injury after MI

To assess the contribution of CD8^+^ T cells to post-MI injury, *Pd1*^*−/−*^*Ctla4*^*+/−*^ mice were treated with an anti-CD8α monoclonal antibody to deplete this subset, and cardiac outcomes were evaluated at 4, 7, and 28 days after infarction (**Figure. 3A; Figure S5A**). CD8^+^ T-cell depletion significantly improved post-MI survival (**Figure 3B**). Masson’s trichrome staining revealed a smaller infarct size in depleted mice (**Figure 3C**). Both TUNEL staining and immunoblotting for cleaved caspase-3 showed reduced cardiomyocyte apoptosis, particularly in the border zone, following depletion (**Figure 3D; Figure S5B**). Echocardiography demonstrated improved cardiac function in anti-CD8α-treated mice, evidenced by increased left ventricular ejection fraction and fractional shortening, along with attenuated chamber dilation (**Figure 3E and 3F; Table S2**). Furthermore, depletion attenuated pathological remodeling: the heart-weight-to-body-weight ratio and cardiomyocyte size detected by WGA staining were reduced, indicating less hypertrophy (**Figure 3G-3H**). Consistent with the functional and structural improvements, mRNA levels of key hypertrophy markers—atrial natriuretic peptide (*Nppa*), brain natriuretic peptide (*Nppb*), and myosin heavy chain 7 (*Myh7*)—were significantly reduced in the hearts of CD8^+^ T-cell-depleted mice (**Figure S5C**). Furthermore, CD8^+^ T-cell depletion ameliorated interstitial fibrosis, as evidenced by histology and a concomitant decrease in the expression of the canonical fibrotic genes fibronectin 1 (*Fn1*), collagen type I alpha 1 (*Col1a1*), and transforming growth factor beta (*Tgfb*) (**Figure S5D and S5E**).In summary, CD8^+^ T-cell depletion improved survival, cardiac function, and attenuated apoptosis, hypertrophy, and fibrosis in *Pd1*^*−/−*^*Ctla4*^*+/−*^ mice after MI.

**Figure 3:**
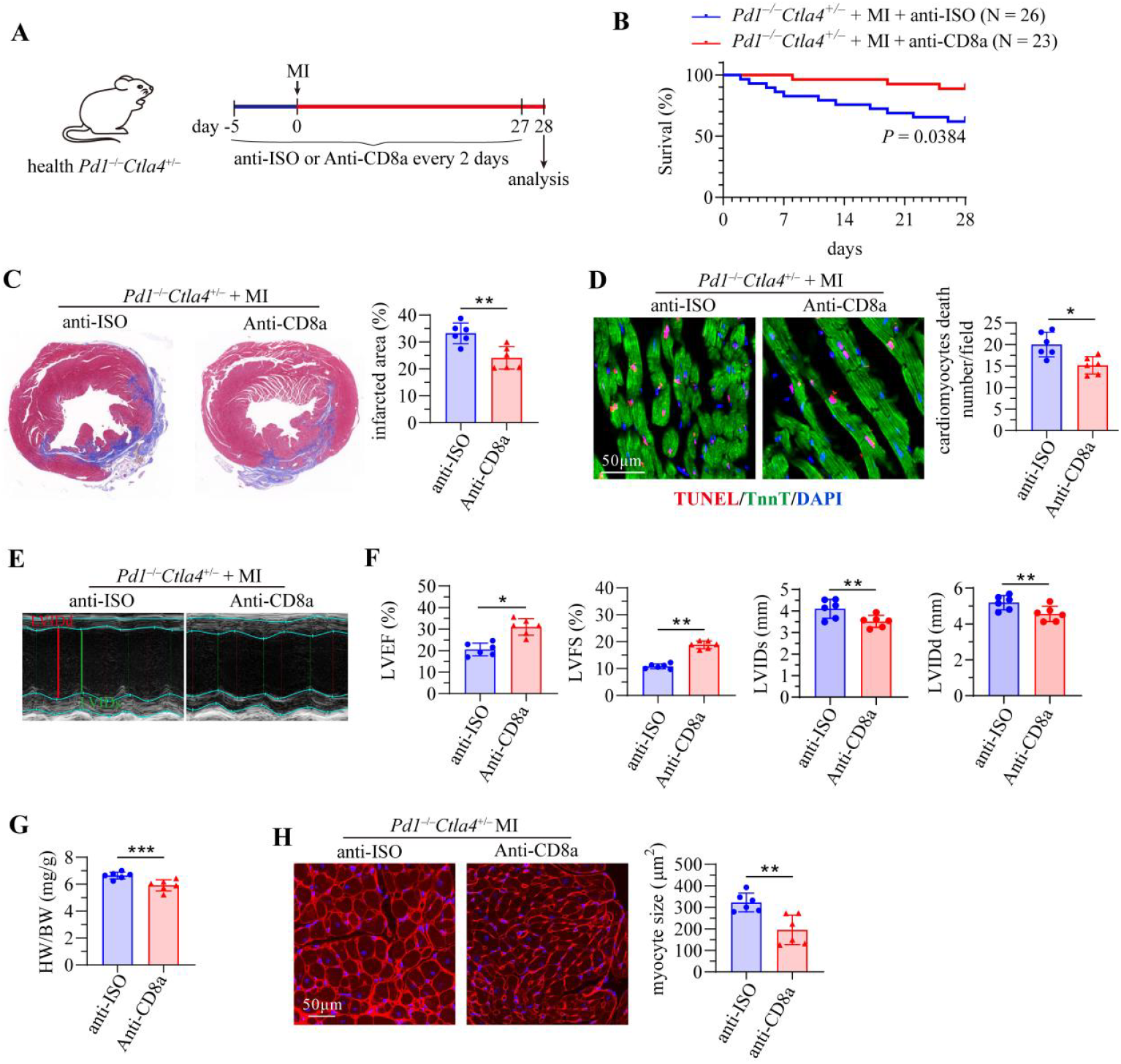
CD8^+^ T-cell depletion mitigates post-MI cardiac injury. **A**, Schematic of the experimental timeline. 12-week-old asymptomatic *Pd1*^*−/−*^*Ctla4*^*+/−*^ mice received intraperitoneal injections of either anti-CD8α antibody or anti-ISO type control antibody every 2 days, starting 5 days before and continuing until 28 days after MI. **B**, Kaplan–Meier survival curves over 4 weeks post-MI. **C**, Representative images of Terminal deoxynucleotidyl transferase dUTP Nick-End Labeling (TUNEL) staining (left) and quantification of TUNEL^+^ cardiomyocytes (right) at 28 days post-MI. Cardiomyocytes are stained green (cTnT), apoptotic nuclei red (TUNEL), and all nuclei blue (DAPI). **D**, Representative TUNEL staining images and quantification of apoptotic (TUNEL^+^) cardiomyocytes in the peri-infarct border zone 28 days after surgery. TUNEL^+^ nuclei are shown in red, cTnT^+^ cardiomyocytes in green, and DAPI-counterstained nuclei in blue. **E**, Representative M-mode echocardiographic tracings. **F**, Quantification of left ventricular ejection fraction (LVEF), fractional shortening (FS), and LV internal dimensions in systole (LVIDs) and diastole (LVIDd) by echocardiography. **G**, Quantification of heart weight-to-body weight (HW/BW) ratio. **H**, Representative wheat germ agglutinin (WGA) staining images (left) and quantification of cardiomyocyte cross-sectional area (right). Data are presented as mean±SD. ns, not significant; \**P*< 0.05, \*\**P*< 0.01, \*\*\**P*< 0.001.

### Depletion of CD8^+^ T cells attenuates macrophage inflammation

To investigate the role of expanded CD8^+^ T cells in regulating macrophage-driven inflammation after myocardial infarction (MI), *Pd1*^*−/−*^*Ctla4*^*+/−*^ mice were treated with an anti-CD8α monoclonal antibody to deplete CD8^+^ T cells (**Figure 4A**). Flow cytometry confirmed efficient depletion of CD8^+^ T cells without affecting the number of CD64^+^ macrophages in the heart (**Suppl. Figure S4A and S4B**). Depletion of CD8^+^ T cells led to decreased cardiac mRNA levels of interferon-γ (*Ifng*) and granzyme B (*Gzmb*), reflecting reduced T-cell derived inflammatory activity. Concurrently, mRNA levels of the macrophage-associated cytokines interleukin-1β (*Il1b*), interleukin-6 (*Il6*), tumor necrosis factor-α *Tnfa*, and C-C chemokine ligand 2 (*Ccl2*) were also diminished, indicating attenuated macrophage inflammation (**Figure 4B**). This reduction in inflammatory activation was further validated in flow-sorted macrophages, which showed lower expression of *Il1b, Il6*, and *Tnf*a following CD8^+^ T-cell depletion (**Figure 4C**). Moreover, flow cytometric analysis revealed decreased CD86 and increased CD206 expression on macrophages in the hearts of CD8^+^ T-cell-depleted mice, suggesting a shift toward a less inflammatory phenotype (**Figure 4D**). Together, these data indicate that CD8^+^ T cells exacerbate early post-MI cardiac inflammation by promoting a pro-inflammatory macrophage state, and their depletion alleviates this response.

**Figure 4:**
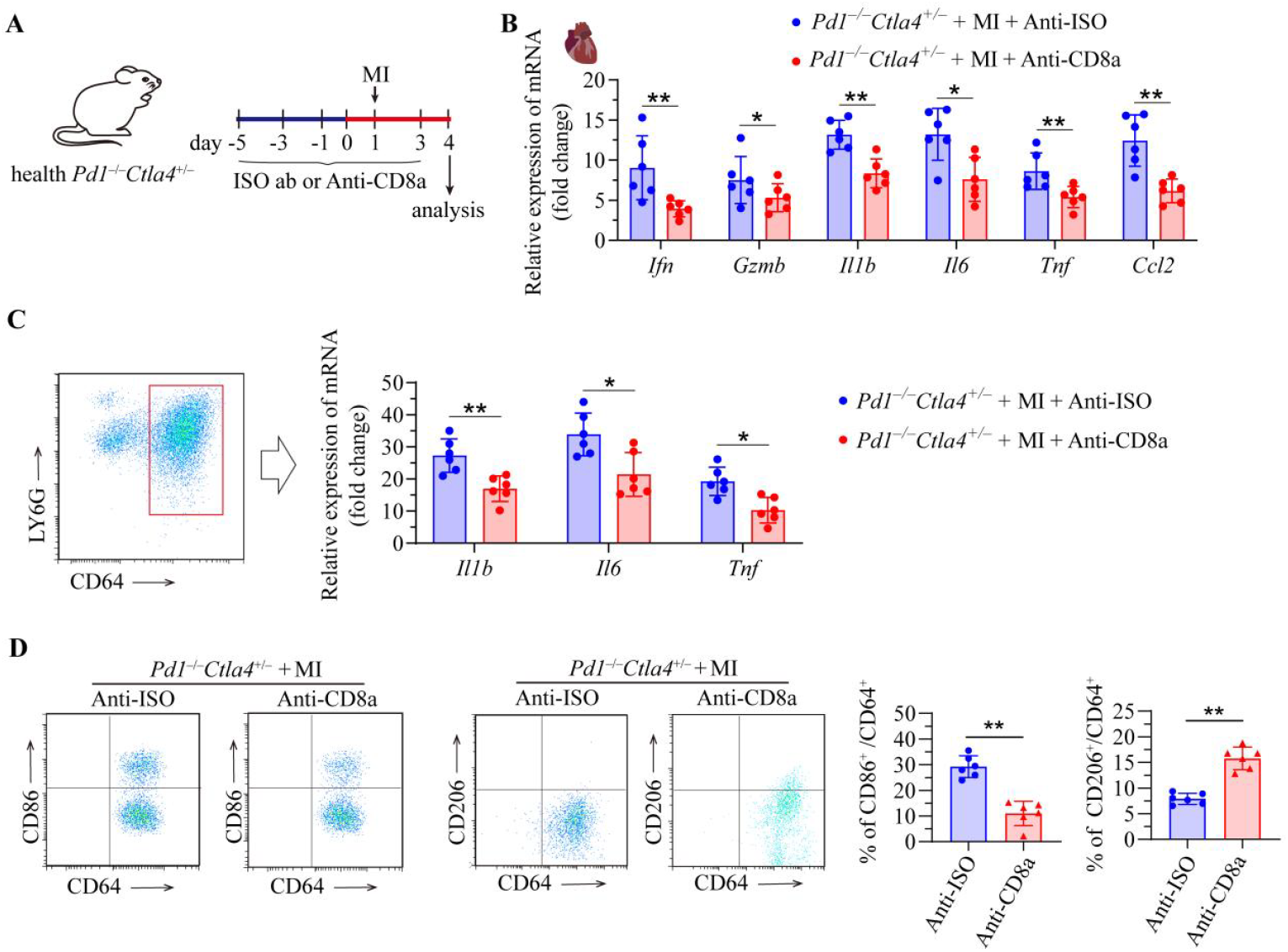
Depletion of CD8^+^ T Cells attenuate macrophage inflammation. **A**, Schematic of the experimental timeline. 12-week-old asymptomatic *Pd1*^−/−^*Ctla4*^+/−^ mice received intraperitoneal injections of either anti-CD8α antibody or isotype control antibody every 2 days before and after MI surgery. Heart tissues were harvested for analysis 4 days post-MI. **B**, Quantitative PCR analysis of mRNA expression levels of *Il1b, Il6, Tnfa*, and *Ccl2* in the hearts of *Pd1*^−/−^*Ctla4*^+/−^ mice. **C**, Strategy of flow cytometry sorting CD64^+^ macrophages from hearts of *Pd1*^−/−^*Ctla4*^+/−^ mice. **D**, Quantitative PCR analysis of mRNA expression levels of *Ifng, GzmB, Il1b, Il6* and *Tnfa* in the macrophages sorted from hearts of *Pd1*^−/−^*Ctla4*^+/−^ mice. **E**, Representative flow cytometric images and corresponding quantitative analysis of CD86 and CD206 expressions in CD64^+^ macrophages. Data are presented as mean±SD. ns, not significant; \**P*< 0.05, \*\**P*< 0.01, \*\*\**P*< 0.001.

### CD8^+^ T Cell-Derived IFN-γ Activates the JAK-STAT1 Pathway in the Infarcted Heart

To characterize the inflammatory properties of cardiac CD8^+^ T cells post-MI, we performed MI surgery on wild-type (*Pd1*^+/+^*Ctla4*^+/+^) and *Pd1*^−/−^*Ctla4*^+/−^ mice. Heart tissues were collected at 4 or 7 days post-MI for analysis (**Figure 5A**). CD8^+^ T cells were sorted by flow cytometry for quantitative PCR analysis (**Figure 5B**). Compared to wild-type, cardiac CD8^+^ T cells from *Pd1*^−/−^*Ctla4*^+/−^ mice showed elevated mRNA levels of both *Ifng* and *GzmB*. The increase in *Ifng* was particularly pronounced, indicating that it is the dominantly induced effector molecule and pointing to enhanced inflammatory T cell function (**Figure 5C**). Immunofluorescence staining confirmed elevated IFN-γ protein expression specifically in these CD8^+^ T cells (**Figure 5D**). Correspondingly, western blot analysis demonstrated an increased overall level of IFN-γ protein in the heart tissue of *Pd1*^−/−^*Ctla4*^+/−^ mice (**Figure 5E**).

**Figure 5:**
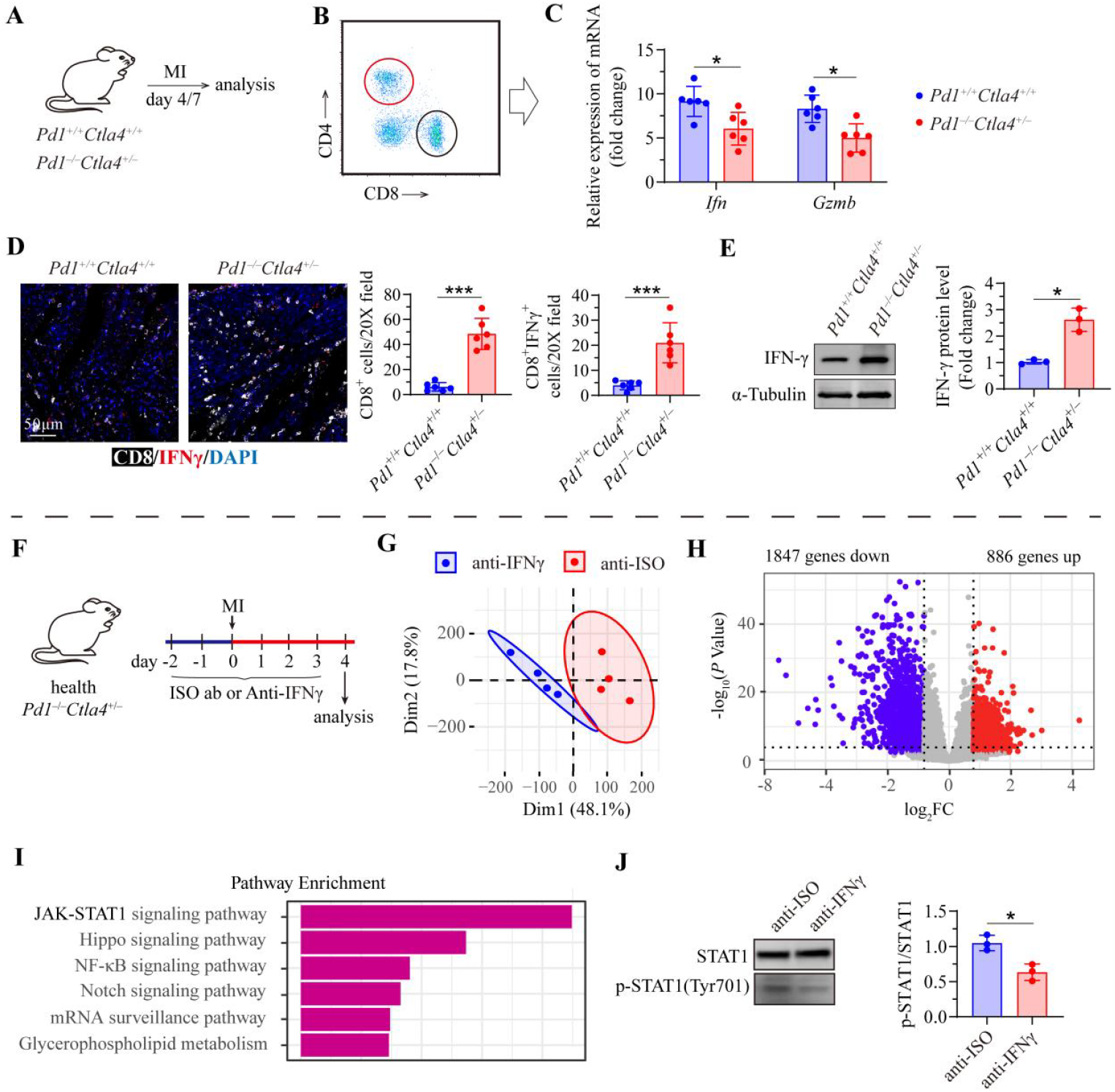
Deficiency of *Pd1* and *Ctla4* enhances activation of the JAK-STAT1 signaling pathway after MI. **A**, Schematic of the experimental timeline. 12-week-old wild-type (*Pd1*^*+/+*^*Ctla4*^*+/+*^) and *Pd1*^*−/−*^*Ctla4*^*+/−*^ mice underwent MI surgery, with analysis performed 4 days later. **B**, Principal component analysis (PCA) of myocardial gene expression profiles, showing segregation between the *Pd1*^*+/+*^*Ctla4*^*+/+*^ + MI group (blue) and the *Pd1*^*−/−*^*Ctla4*^*+/−*^ + MI group (red). **C**, Volcano plot of differentially expressed genes in *Pd1*^*+/+*^*Ctla4*^*+/+*^ + MI *versus Pd1*^*−/−*^*Ctla4*^*+/−*^ + MI hearts. Red and blue dots denote upregulated and downregulated genes, respectively (*P* < 0.05, fold change ≥ 1.5). **D**, Heatmap of the top 10 upregulated and downregulated differentially expressed genes between the two groups. **E**, Kyoto Encyclopedia of Genes and Genomes (KEGG) pathway enrichment analysis of upregulated pathways in *Pd1*^*−/−*^*Ctla4*^*+/−*^ + MI hearts. **F**, Schematic of the experimental timeline. 12-week-old asymptomatic *Pd1*^*−/−*^*Ctla4*^*+/−*^ mice received daily intraperitoneal injections of either anti-IFNγ antibody or isotype control antibody, starting 2 days before and continuing until 4 days after MI surgery. Heart tissue was collected 4 days post - MI for analysis. **G**, Principal component analysis (PCA) of transcriptomic profiles from anti-IFNγ-treated and control hearts. **H**, Volcano plot of differentially expressed genes in anti-IFNγ-treated *versus* control hearts. **I**, KEGG pathway enrichment analysis of genes downregulated following anti-IFNγ treatment. **J**, Representative western blots of STAT1 phosphorylated at Tyr701 (p-STAT1) and total STAT1 (left) and quantitative analysis of the p-STAT1/STAT1 ratio (right) in myocardial tissue. Data are presented as mean±SD. \**P*<0.05, \*\**P*<0.01.

To define the functional role of this elevated IFNγ, *Pd1*^−/−^*Ctla4*^+/−^ mice undergoing MI were treated with an IFN-γ-neutralizing antibody (**Figure 5F**). RNA sequencing of heart tissue revealed that IFN-γ blockade markedly altered the cardiac transcriptome. Principal component analysis (PCA) showed a clear separation in global gene expression profiles between anti-IFN-γ-treated and control groups (**Figure 5G**). A volcano plot identified approximately 1,000 downregulated and 500 upregulated genes upon IFNγ neutralization (**Figure 5H**). KEGG pathway enrichment analysis of the downregulated genes indicated significant suppression of the JAK-STAT pathway, particularly involving STAT1 (**Figure 5I**). A circular interaction network further illustrated the inhibition centered on IFNGR1 and downstream JAK-STAT components (**Figure 7A and 7B**). Finally, western blot analysis confirmed that IFNγ blockade strongly inhibited the phosphorylation (activation) of STAT1 in heart tissue (**Figure 5J**).

Collectively, these results demonstrate that PD-1/CTLA-4 deficiency potentiates CD8^+^ T cell effector function in the infarcted heart, characterized by robust IFN-γ production. Furthermore, this IFN-γ drives pathogenic inflammatory signaling within the heart primarily through activation of the JAK-STAT1 pathway.

### JAK-STAT1 Inhibition Attenuates Macrophage and Myocardial Inflammation

Given the robust activation of the JAK-STAT1 pathway in *Pd1*^−/−^*Ctla4*^+/−^ hearts after MI, we asked whether pharmacologically inhibiting this axis would attenuate macrophage and myocardial inflammation. *Pd1*^−/−^*Ctla4*^+/−^ mice were treated with the JAK inhibitor Tofacitinib (**Figure 6A; Figure S8A**). Western blot analysis confirmed that Tofacitinib effectively suppressed STAT1 phosphorylation in the heart (**Figure 8B**). Both mRNA and protein levels of inflammatory mediators, including *Il1b, Il6, Tnf, Ccl2, Ccl3*, and *Ccl4*, were reduced in Tofacitinib-treated hearts. Flow-sorted cardiac macrophages from these mice also showed decreased expression of inflammatory genes such as *Ifng, Il1b, Il6, Tnfa*, and *Ccl2* (**Figure 6B**).

**Figure 6:**
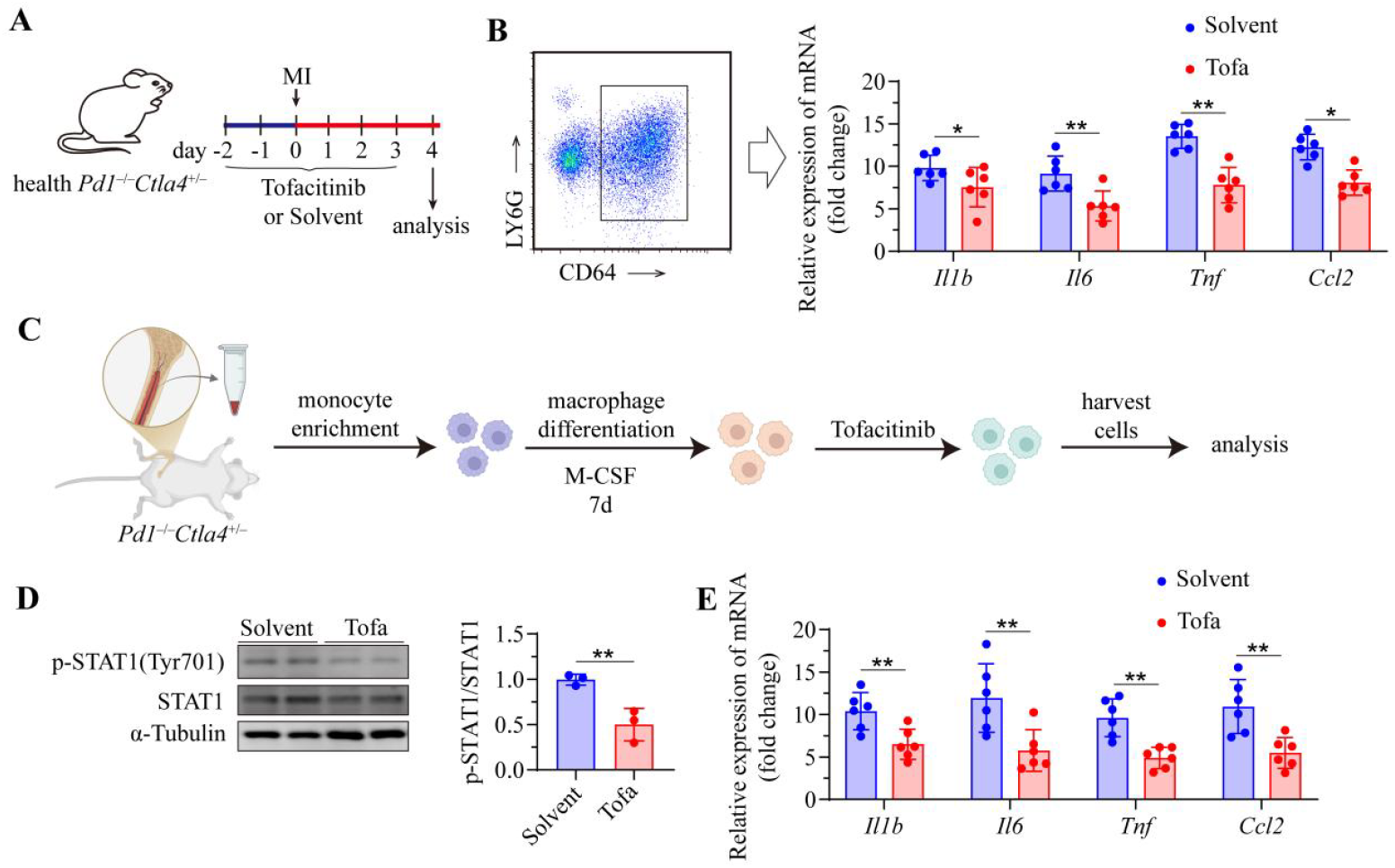
JAK-STAT1 Inhibition attenuates macrophage inflammation. **A**, Schematic of the experimental timeline. 12-week-old asymptomatic *Pd1*^−/−^*Ctla4*^+/−^ mice received intraperitoneal injections of either anti-IFNγ antibody or isotype control antibody every 2 days before and after MI surgery. Heart tissues were harvested for analysis 4 days post-MI. **B**, Cardiac CD64^+^ macrophages sorting strategy and quantitative PCR analysis of mRNA expression levels of *Ifng, GzmB, Il6, Tnfa, Il1b*, and *Ccl2* in the hearts of MI-injured *Pd1*^−/−^*Ctla4*^+/−^ mice treated with Tofacitinib and solvent. **C**, Schematic diagram of bone marrow-derived macrophage (BMDM) isolation from *Pd1*^−/−^*Ctla4*^+/−^ mice and the following culture procedure. **D**, Representative western blots of myocardial p-STAT1 (Tyr701) and total STAT1, and quantitative analysis of protein levels. **E**, Quantitative PCR analysis of mRNA expression levels of *Ifng, GzmB, Il6, Tnfa, Il1b*, and *Ccl2* in BMDMs treated with Tofacitinib and solvent. Data are presented as mean±SD. ns, not significant; \**P* < 0.05, \*\**P*< 0.01, \*\*\**P*< 0.001.

**Figure 7:**
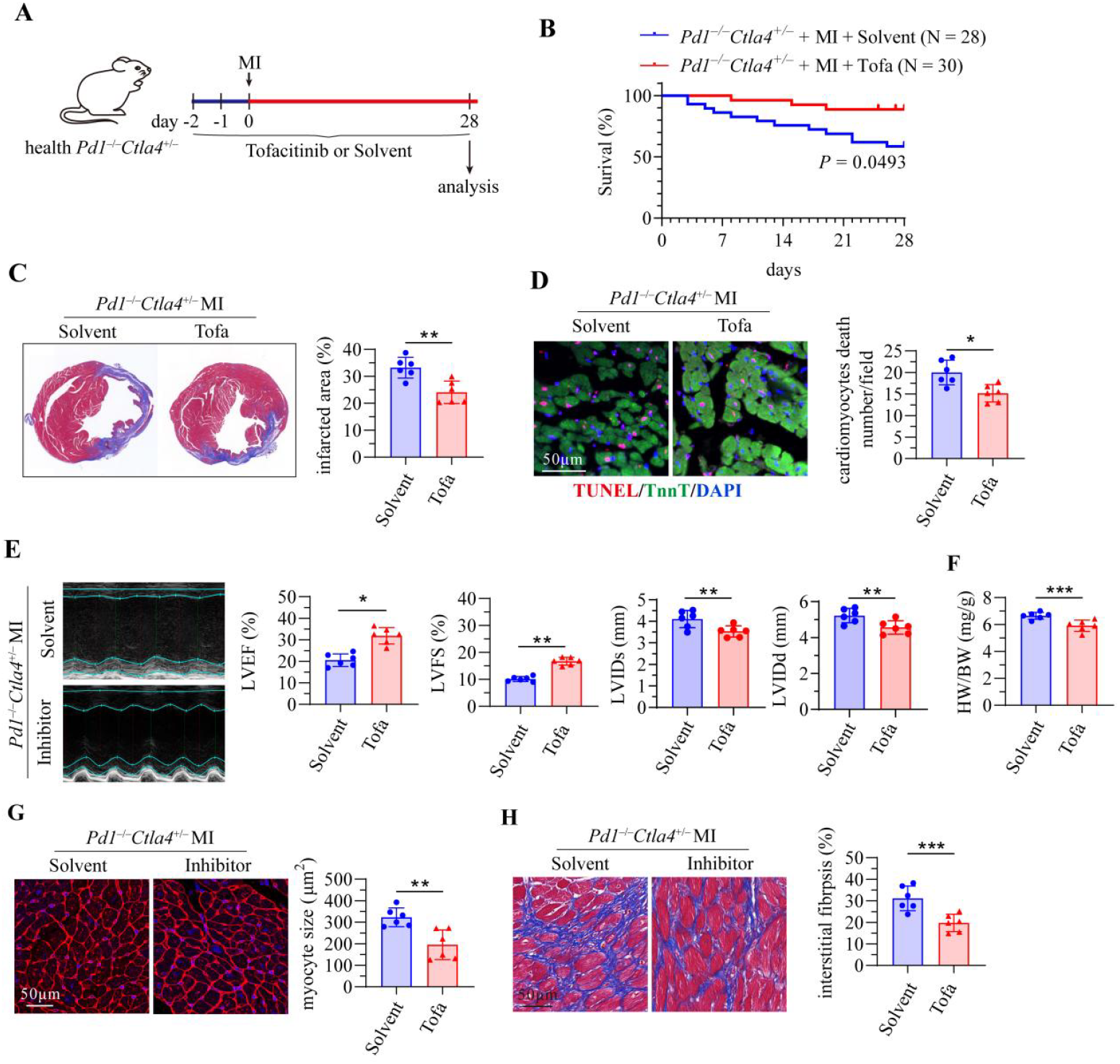
JAK-STAT1 inhibition improves cardiac function and remodeling after MI. **A**, Schematic of the experimental timeline. 12-week-old asymptomatic *Pd1*^−/−^*Ctla4*^+/−^ mice received daily oral gavage of the JAK inhibitor Tofacitinib or vehicle, starting 2 days before MI and continuing every 2 days until 28 days after surgery. Echocardiography was performed before tissue collection at the endpoint. **B**, Kaplan–Meier survival curves over 4 weeks post-MI. **C**, Representative Masson’s trichrome-stained cross-sections of the left ventricle (left) and quantification of infarct size as a percentage of the left ventricular area (right). **D**, Representative TUNEL staining images and quantification of apoptotic (TUNEL^+^) cardiomyocytes in the peri-infarct border zone 28 days after surgery. TUNEL^+^ nuclei are shown in red, cTnT^+^ cardiomyocytes in green, and DAPI-counterstained nuclei in blue. **E**, Representative M-mode echocardiographic tracings (left) and quantification of left ventricular ejection fraction (LVEF), fractional shortening (LVFS), and left ventricular internal dimensions in diastole (LVIDd) and systole (LVIDs) (right). **F**, Heart-weight-to-body-weight (HW/BW) ratio. **G**, Representative wheat germ agglutinin (WGA) staining images (left) and quantification of cardiomyocyte cross-sectional area (right). **H**, Representative Masson’s trichrome staining of the peri-infarct border zone (left) and quantification of interstitial fibrosis area (right). Data are presented as mean±SD. ns indicates non-significance, \**P*<0.05, *P*<0.01,\**P*<0.001.

In parallel, bone marrow-derived macrophages (BMDMs) from *Pd1*^−/−^*Ctla4*^+/−^ mice were exposed to Tofacitinib in vitro (**Figure 6C**). The inhibitor markedly diminished STAT1 phosphorylation at Tyr701 and lowered total STAT1 protein levels (**Figure 6D**). Consistent with this, qPCR analysis demonstrated downregulation of key inflammatory cytokines in BMDMs (**Figure 6E**). Together, these findings indicate that JAK-STAT1 inhibition suppresses both macrophage-driven inflammation and global myocardial inflammation after MI.

### Pharmacologic Inhibition of JAK-STAT1 Improves Cardiac Function and Remodeling After MI injury

*Pd1*^−/−^*Ctla4*^+/−^ mice subjected to MI were treated with the JAK inhibitor Tofacitinib or vehicle and analyzed at 4, 7, or 28 days post-injury (**Figure 7A; Figure S9A**). Tofacitinib treatment significantly improved post-MI survival (**Figure 7B**) and reduced infarct size (**Figure 7C**). Both TUNEL staining and western blot analysis demonstrated attenuated cardiomyocyte apoptosis in inhibitor-treated hearts (**Figure 7D; Figure S9B**). Echocardiography revealed substantially improved cardiac function, evidenced by increased left ventricular ejection fraction (LVEF) and fractional shortening (LVFS), as well as attenuated ventricular dilation (reduced LVIDd and LVIDs) (**Figure 7E; Table S3**). Furthermore, Tofacitinib ameliorated pathological remodeling. Cardiac hypertrophy was reduced, as shown by a lower heart weight to body weight ratio, decreased cardiomyocyte cross-sectional area (wheat germ agglutinin staining), and downregulated mRNA expression of hypertrophy markers—atrial natriuretic peptide (*Nppa*), brain natriuretic peptide (*Nppb*), and myosin heavy chain 7 (*Myh7*) (**Figure 7F and 7G; Figure S9C**). Interstitial fibrosis was also attenuated, indicated by reduced collagen deposition (Masson’s trichrome staining) and decreased mRNA levels of fibrotic genes, including fibronectin 1 (*Fn1*), collagen type I alpha 1 chain (*Col1a1*), and transforming growth factor beta (*Tgfb*) (**Figure 7H; Figure S9D**). Collectively, pharmacological inhibition of the JAK-STAT1 axis improved survival, restored cardiac function, and mitigated key features of adverse post-MI remodeling in *Pd1*^−/−^*Ctla4*^+/−^ mice.

## Disscussion

A critical unresolved question in cardio-oncology is whether immune checkpoint inhibitors (ICIs), even in the absence of overt myocarditis, predispose the heart to more severe injury during an acute ischemic event^[44–46]^. This study provides direct experimental evidence supporting this concerning possibility. Our data indicate that genetic mimicry of combined PD-1/CTLA-4 blockade predisposes the heart to a latent inflammatory state. While not sufficient to cause spontaneous myocarditis in most mice, this pre-conditioning led to a dramatically exaggerated inflammatory response, worse cardiac dysfunction, and significantly higher mortality following myocardial infarction (MI). This translates to a critical clinical implication: an acute MI in a patient receiving combination ICI therapy—even one without prior cardiac symptoms—may trigger a disproportionately severe inflammatory storm and result in worse outcomes. Thus, acute infarction may represent another serious, context-dependent manifestation of ICI-related cardiotoxicity beyond classic myocarditis^[47, 48]^.

Mechanistically, we dissect a novel pathogenic axis wherein combined checkpoint deficiency exacerbates post-MI injury. The aggravation was driven by a pronounced infiltration of activated, interferon-γ (IFN-γ)-producing CD8^+^ T cells. These cells were essential drivers of pathology, as their depletion attenuated global injury and, specifically, mitigated macrophage-driven inflammation. The CD8^+^ T cell-derived IFN-γ activated the JAK-STAT1 signaling pathway in macrophages, polarizing them toward a pro-inflammatory state. Pharmacological inhibition of this pathway with Tofacitinib successfully suppressed inflammation and ameliorated all aspects of post-MI injury. The central conclusion is that combined checkpoint inhibition worsens post-MI outcomes by promoting a CD8^+^ T cell-dependent activation of the macrophage JAK-STAT1 pathway.

The expansion of pathogenic CD8^+^ T cells can be attributed to a “perfect storm” of antigen exposure and lost inhibition. MI releases cardiac antigens, creating a pro-inflammatory milieu. PD-1 and CTLA-4 normally act as crucial brakes on T-cell activation. Their combined genetic removal (or clinical blockade) releases these brakes, leading to hyper-responsive CD8^+^ T cells with enhanced clonal expansion, survival, and effector function, as evidenced by elevated IFN-γ and granzyme B^[7, 49, 50]^.

Our study points directly to two promising therapeutic strategies for mitigating cardiac injury in ICI-treated patients suffering an MI: targeting the pathogenic CD8^+^ T cells or the downstream JAK-STAT1 pathway in macrophages. Short-term, targeted intervention with a JAK inhibitor like Tofacitinib in the acute phase of MI could be a translatable adjunctive therapy to dampen the exaggerated immune response without completely negating the anti-tumor benefits of ICIs.

This study has certain limitations. First, we employed a genetic knockout model to simulate the chronic systemic effects of combined ICI therapy^[51, 52]^. While this effectively recapitulates a state of checkpoint deficiency, it does not fully mirror the pharmacokinetics of administering monoclonal antibodies to wild-type animals or patients^[53–55]^. Future studies using anti-PD-1/CTLA-4 antibodies in wild-type or humanized mouse models would strengthen clinical translatability. Second, our investigation focused primarily on the early and subacute phase post-MI^[56–58]^. The long-term consequences of this primed inflammatory state on chronic cardiac remodeling, heart failure development, and the potential interplay with atherosclerosis warrant further exploration^[59–61]^.

Beyond validating the CD8^+^ T cell-macrophage-STAT1 axis, future research should elucidate the precise cardiac antigens triggering this aberrant T-cell response and investigate the role of other immune cells, such as dendritic cells in antigen presentation or Tregs in failed immunoregulation^[62–65]^. Furthermore, identifying biomarkers (e.g., elevated IFN-γ or specific T-cell clones) that predict high risk for exaggerated post-ischemic injury in ICI patients could enable personalized monitoring and pre-emptive therapy^[66–68]^.

In summary, this work elucidates a novel mechanism by which combined ICI therapy exacerbates ischemic heart injury, shifting the paradigm to consider MI as a context-dependent immune-related adverse event. It provides a strong scientific rationale for vigilant cardiac monitoring in ICI recipients and offers tangible therapeutic targets, paving the way for developing cardioprotective strategies that could improve the overall survivorship of cancer patients.

## Nonstandard Abbreviations and Acronyms

ICIs: Immune Checkpoint Inhibitors
MI: Myocardial Infarction
Pd1: Programmed Cell Death Protein 1
Ctla4: Cytotoxic T-Lymphocyte-Associated Protein 4
IFN-γ: Interferon-gamma
Tofa: Tofacitinib

